# Incorporating uncertainty within dynamic interoceptive learning

**DOI:** 10.1101/2023.05.19.538717

**Authors:** Katja Brand, Toby Wise, Alexander J. Hess, Bruce R. Russell, Klaas E. Stephan, Olivia K. Harrison

## Abstract

Interoception, the perception of the internal state of the body, has been shown to be closely linked to emotions and mental health. Of particular interest are interoceptive learning processes that capture associations between environmental cues and body signals as a basis for making homeostatically relevant predictions about the future. Here we extended an interoceptive Breathing Learning Task (BLT) to incorporate continuous measures of prediction certainty, and tested its application using a Rescorla Wagner (RW) associative learning model. Sixteen healthy participants completed the continuous version of the BLT, where they were asked to predict the likelihood of breathing resistances. The task was modified from a previous version and required continuous, rather than binary predictions, in order to include a more precise measure of prediction certainty. The RW model was used to fit a learning rate to each participant’s continuous and binarised predictions, and was additionally extended to test whether learning rates differed according to stimuli valence. The empirical task data demonstrated excellent replicability compared to previously collected data using binary predictions, and the continuous model fits closely captured participant behaviour at the group level. The model extension to estimate different learning rates for negative (i.e. breathing resistance) and positive (i.e. no breathing resistance) trials indicated that learning rates did not significantly differ according to stimuli nature. Furthermore, examining the relationship between estimates of prediction certainty and learning rates with interoceptive and mental health questionnaires demonstrated that fatigue severity was related to both prediction certainty and learning rate, and anxiety sensitivity was related to prediction certainty. The updated task and model show promise for future investigations into interoceptive learning and potential links to mental health.

## 1 INTRODUCTION

Perception goes beyond the path of registration of sensations but involves the active interpretation of sensory inputs. This interpretation is shaped by numerous cognitive factors, such as prior knowledge or expectations, as well as attention. While exteroception refers to the perception of the external environment through the traditional senses of touch, sight, hearing, taste and smell, interoception refers to the perception of the internal state of the body (Khalsa et al., 2018). Interoception includes both conscious and subconscious processes; investigating these processes is critical for understanding brain-body interactions. While one of the main roles of interoception is to drive actions to maintain homeostasis (Petzschner et al., 2017; Pezzulo et al., 2015; Quadt et al., 2018; Stephan et al., 2016), interoception is also thought to play an important role in emotional regulation (Barrett et al., 2004; Critchley et al., 2004; Füstös et al., 2013). Importantly, interoceptive dysfunction has been implicated in a range of psychological disorders such as depression, anxiety and eating disorders (Khalsa et al., 2018; Paulus and Stein, 2010), and is a rapidly expanding field of research (Brewer et al., 2021).

Interoceptive learning refers to the updating of interoceptive predictions. Computational theories of perception suggest that the brain acts as an “inference machine” that continuously updates probabilistic representations (beliefs) about the state of the world (Friston, 2005; Rao and Ballard, 1999) which, in turn, instantiate predictions about future incoming sensory stimuli. By minimising the difference between actual and predicted stimuli (the prediction error), the brain can update its beliefs according to Bayesian principles (Friston, 2005). Numerous experimental studies have provided empirical evidence that exteroceptive processes such as sight and hearing operate in this manner (Chennu et al., 2013; Kok and de Lange, 2014; Lieder et al., 2013; Stefanics et al., 2018), and these theories are now being extended to interoception in order to explain how the brain uses interoceptive stimuli to create a predictive model of the internal state (Barrett and Simmons, 2015; Critchley and Garfinkel, 2017; Gu et al., 2013; Pezzulo et al., 2015; Seth et al., 2012).

Altered interoceptive learning has been proposed to underpin aspects of psychopathology. Paulus et al. (2019) hypothesise that depression and anxiety are potentially linked to two main interoceptive dysfunctions: overly strong expectations, which shape the processing of interoceptive stimuli; and difficulty in updating predictions to reflect changes in the external or internal state, which may involve faulty prediction error signaling. Electroencephalography (EEG) data has shown that individuals with anxiety have altered neural activity when processing predictions and prediction errors in a visual reward-learning task (Hein and Ruiz, 2022), lending support to this theory. Additionally, previous work by Harrison et al. (2021) developed an interoceptive learning task that was paired with functional magnetic resonance imaging (fMRI), using inspiratory resistances as an interoceptive stimulus (Rieger et al., 2020; Frässle et al., 2021). This task was used to investigate the link between anxiety and interoceptive learning regarding breathing stimuli, finding that more anxious individuals showed altered activity in the anterior insula related to prediction certainty compared to less anxious individuals (Harrison et al., 2021). In this study, binary measures of interoceptive predictions were fitted using a simple associative learning model (a Rescorla Wagner model; (Rescorla et al., 1972)), to determine a learning rate for each participant as well as the corresponding trajectories for predictions and prediction errors.

The current project builds on this interoceptive breathing learning task (BLT) (Harrison et al., 2021) by incorporating direct measures of prediction certainty in place of binary predictions. Past research has suggested that mental health disorders such as anxiety may be associated with an altered response to uncertainty (Grupe and Nitschke, 2013). Therefore, to more accurately capture measures of (un)certainty, the BLT from Harrison et al. (2021) was modified to elicit continuous rather than binary prediction data, thus incorporating a direct measure of certainty surrounding predictions. Additionally, there is evidence that individuals learn more quickly (i.e. adapt their predictions more rapidly) in response to negative outcomes (Aylward et al., 2019; Khdour et al., 2016). Therefore, a model extension that incorporated stimuli valence (i.e. the presence of a breathing resistance [negative stimulus] or the absence of a breathing resistance [positive stimulus]) was also employed to examine whether a single or two separate learning rates would better explain the behavioural data. The modified BLT and learning model were then tested on a sample of 16 healthy participants, and the results were compared to the data collected by Harrison et al. (2021). Additionally, the estimated interoceptive learning parameters were compared to measures of anxiety, depression, affect and subjective interoception in an exploratory analysis.

## 2 MATERIALS AND EQUIPMENT

### 2.1 BLT equipment setup

In order to deliver the breathing resistances to participants during the BLT, an inspiratory resistance circuit was utilised (Rieger et al., 2020). Participants were fitted with a silicone facemask that was adjusted to make a tight seal around mouth and nose. This was then connected to a single-use bacterial and viral filter within the circuit, and inspiratory resistances were induced via an automatically controlled solenoid valve. This valve allowed air to be drawn into the circuit either directly from the environment (with no resistance) or through a PowerBreathe device, which was set to deliver the predetermined amount of resistance (30% of a participant’s maximal inspiratory pressure). The circuit also included a pressure sampling line and spirometer, allowing for continuous measurements of inspiratory pressure and flow to be taken throughout the task. For a diagram of the full circuit used see Figure 1.

**Figure 1.**
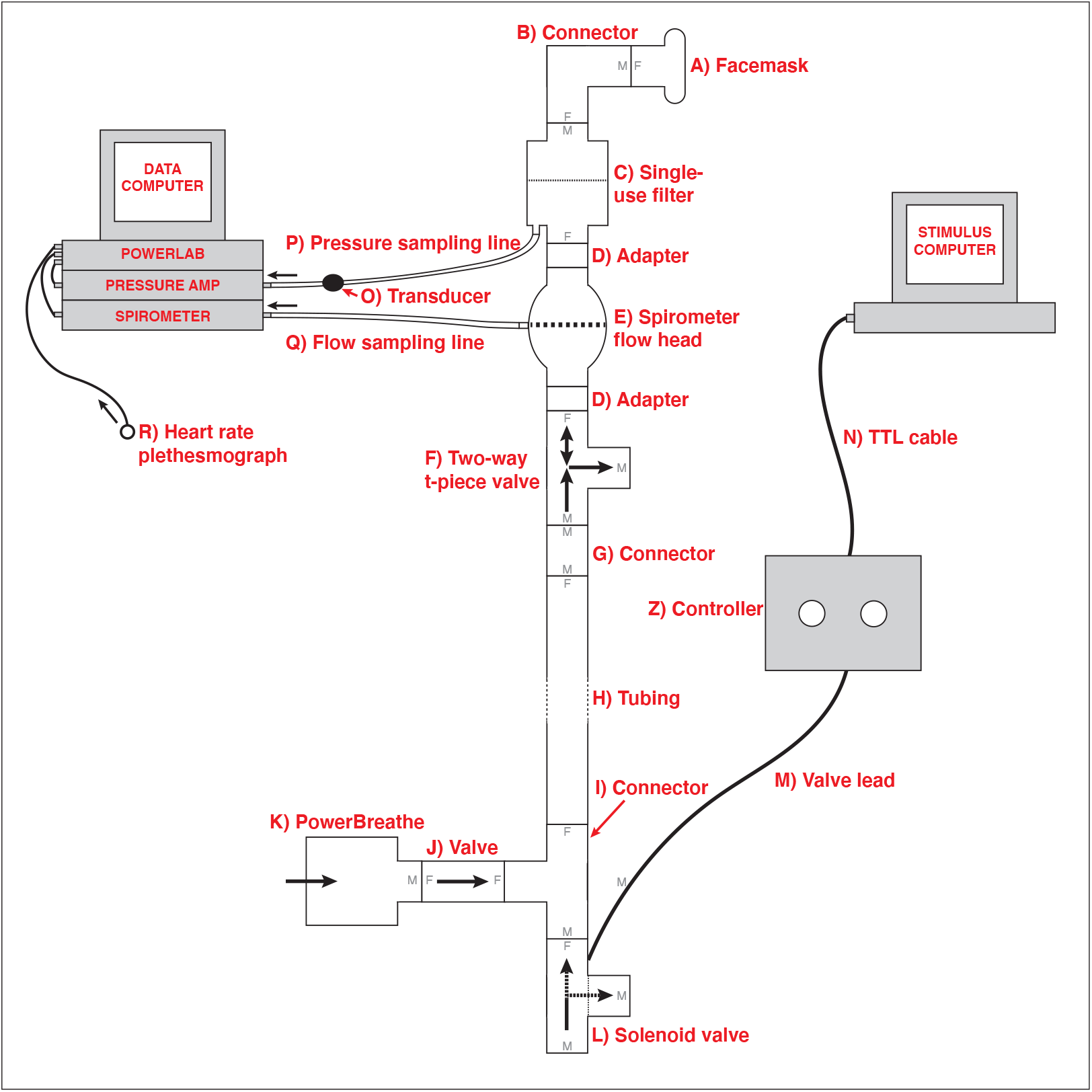
Schematic of the inspiratory resistance circuit used. Participants were connected to the circuit via a facemask covering their mouth and nose which was attached to a bacterial and viral filter (labelled C). A sampling line to a pressure transducer (labelled O) and amplifier was used to record inspiratory pressure, and a spirometer (labelled E) was used to measure inspiratory flow. Inspiratory resistances were induced automatically via the stimulus computer, controller box and solenoid valve (labelled L) to redirect unobstructed air from the environment to the PowerBreathe device (labelled K), which provided a set magnitude of resistance. Figure adapted from Rieger et al. (2020) under a CC-BY licence.

## 3 METHODS

### 3.1 Participants and recruitment

In order to test the continuous response version of the BLT, data was gathered from 16 healthy volunteers. Participants were aged 19-42 years (mean age: 23y; 4M, 12F), and were pre-screened according to the following criteria:

- Aged 18-45
- Regularly exercising no more than once per week
- Non-smoker or light smoker (smoking or vaping once per week or less)
- Not on any regular medication at time of study (except the oral contraceptive pill)
- Full colour-vision
- Not suffering from any chronic medical conditions, including current or past history of brain injury or breathing disorder
- No past or current diagnoses of schizophrenia, bipolar disorder, drug addiction, or psychosis
- Not pregnant or breastfeeding

Participants were recruited from the community using study advertisements. All participants signed a written, informed consent, and the study was approved by the New Zealand Health and Disability Ethics Committee (HDEC) (Ethics approval 20/CEN/168).

Data from a separate group of eight participants (4M, 4F) were used to determine model priors. These participants had completed an earlier version of the BLT using binarised responses and their data had initially been used by (Harrison et al., 2021). All participants signed a written, informed consent, and the study was approved by the Cantonal Ethics Committee Zurich (Ethics approval BASEC-No. 2017-02330).

### 3.2 Procedure

Participants who were selected for the study following online pre-screening were asked to complete a series of questionnaires (details in Section 3.2.1) followed by BLT (see Section 3.2.2). Both tasks required 30-45min to complete.

#### 3.2.1 Questionnaires

Following online pre-screening and informed consent, participants were firstly asked to fill in a number of questionnaires on the lab computer. The questionnaires presented to the participants were designed to capture subjective affective measures as well as general and breathing-specific interoceptive beliefs.

Affective qualities that were measured included state anxiety (measured by the Spielberger Trait Anxiety Inventory; STAI-S; Spielberger et al., 1970), symptoms of generalised anxiety disorder (Generalised Anxiety Disorder Questionnaire; GAD-7; Spitzer et al., 2006), anxiety sensitivity (Anxiety Sensitivity Index; ASI-3; Taylor et al., 2007), symptoms of depression (Centre for Epidemiologic Studies Depression Scale; CES-D; Radloff, 1977), as well as general positive and negative affect (Positive Affect Negative Affect Schedule; PANAS; Watson et al., 1988).

Self-reported interoceptive awareness was measured by the Multidimensional Assessment of Interoceptive Awareness Questionnaire (MAIA; Mehling et al., 2012). Two further questionnaires measured the tendency to catastrophise in response to breathlessness (Pain Catastrophising Scale, adapted to replace pain with breathlessness; PCS-B; Sullivan et al., 1995), as well as awareness and vigilance surrounding breathlessness (Pain Vigilance and Awareness Questionnaire, again replacing pain with breathlessness; PVAQ-B; McCracken, 1997), in line with previous research (Harrison et al., 2021; Herigstad et al., 2017).

Additional facets related to mental health were measured by the following questionnaires: General Self Efficacy scale (GSE; Schwarzer et al., 1997) which measured self-efficacy, Connor Davidson Resilience Scale (CD-RISC; Connor and Davidson, 2003) for resilience, and the Fatigue Severity Scale (FSS; Krupp et al., 1989) for fatigue. Finally, trait anxiety was measured by the Spielberger Trait Anxiety Inventory (STAI-T; Spielberger et al., 1970), which participants filled out during the online pre-screening process.

#### 3.2.2 BLT procedure

After filling out the questionnaires, participants completed the BLT. In order to set an appropriate level of breathing resistance for this task, the maximum inspiratory pressure (MIP) was first measured and recorded for each participant using a PowerBreathe device (PowerBreathe International Ltd, Warwickshire, UK). The resistance magnitude for the BLT was then set to 30% of the participant’s MIP. Once the breathing resistance had been calibrated to the participant, they were fitted with a breathing mask (Hans Rudolph, Kansas City, MO, United States) which was connected to the breathing circuit (see Section 2.1 for details), and tested to ensure that they could feel the resistances. In two cases, the participants were unable to perceive the resistances when operating at their normal tidal volume, and here the resistance was increased to 50% of their MIP to accommodate for this. Written instructions for the BLT were given to the participant to read (see Figure S1 in Supplementary Materials) and these were repeated verbally, as well as allowing the participant to ask questions to ensure they understood the task. Participants were also given a practice version of the task with six trials before beginning the actual task.

During the task, participants were asked how certain they were that there would be an inspiratory resistance following the presentation of one of two visual cues (see Figure 2). Participants were told beforehand that one of the cues had an 80% chance of being followed by a resistance, while the other cue had a 20% chance of being followed by a resistance. They were also told that the pairings of the images with the probabilities could swap during the task, so that the cue previously associated with an 80% chance of resistance now predicted a 20% chance of resistance and vice versa. Participants were not informed of which cue started with which probability, or when the switches would occur.

**Figure 2.**
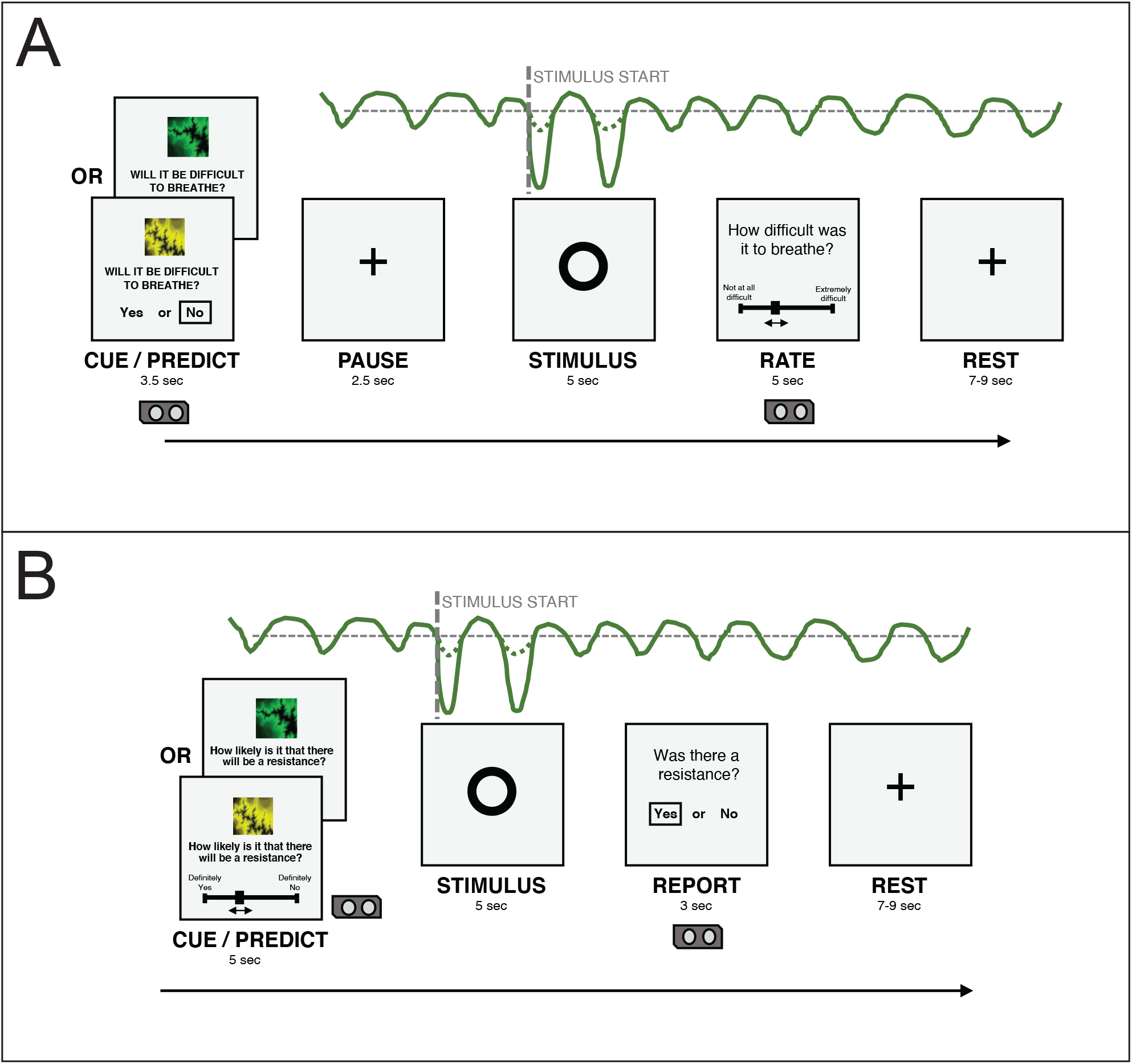
**(A)** An overview of the structure of a single trial in the version of the BLT used in Harrison et al. (2021). **(B)** An overview of the structure of a single trial in the version of the BLT used for the current study. The main changes are that the cue/prediction phase was changed to allow continuous predictions via a slider (with a longer timeframe for this phase to compensate), while the report phase was change to a binary yes/no response to simplify this decision for participants. Additionally the pause before the stimulus was removed after lengthening the cue/prediction phase. Figure adapted from Harrison et al. (2021) under a CC-BY licence.

For each trial, one of the visual cues was displayed on a computer screen for five seconds along with a prompt asking the participant to predict the likelihood of a resistance occurring. Participants’ entered their prediction (along with their certainty in the prediction) by using arrow keys on a keyboard to move a slider on a scale from ‘definitely yes’ to ‘definitely no’. Immediately following this, a circle was shown on the screen for five seconds, during which time the breathing resistance occurred on resistance trials, and no resistance (i.e. normal breathing) occurred on all other trials. Following this period, participants were asked whether or not a resistance had occurred. Participants were then given a rest period of between seven and nine seconds before the next trial began. The protocol for this task was adapted from Harrison et al. (2021) to collect continuous rather than binary prediction data. Figure 2 provides an overview of the structure of each trial and the alterations that were made to gather continuous response data.

In total, the task consisted of 80 trials, which took approximately 30 minutes to complete. During the task, physiological recordings of inspiratory pressure, breathing rate, breathing volume and heart rate were taken using a spirometer and pulse monitor, connected to a PowerLab and recorded using LabChart 8 software (ADInstruments, Dunedin, New Zealand). Participants also wore headphones playing pink noise throughout the task.

The initial pairing of cue-to-resistance was counter-balanced across participants (such that each cue was first paired with an 80% chance of resistance for half of the participants), as well as the position of the ‘definitely yes’ and ‘definitely no’ anchors on the left or right of the screen. The initial pairing was always held constant for the first 30 trials before the pairings were switched (i.e. the cue initially paired with 20% chance of resistance was now paired with 80% chance of resistance and vice versa). The pairings were then switched four more times during the remaining 50 trials at shorter intervals (12-13 trials). The number of trials between each switch was held constant for all participants. This is the same protocol as used and validated in the study by Harrison et al. (2021).

### 3.3 Data processing

Questionnaires were scored according to their respective manuals, with a summary of the relevant scores presented in Section 4.3. For the BLT, data was first checked for missed trials. Data from one participant was excluded from further analysis due to missing more than 10 trials (as predetermined in the analysis plan which can be viewed at https://github.com/IMAGEotago/Katja-BLT-analysisPlan). Next, each participant’s average certainty was determined by taking the absolute value of the difference between their response and 0.5 (with responses being values between 0.0 (definitely no resistance) and 1.0 (definitely resistance) and 0.5 thus representing complete indecision) for each trial and averaging across all trials. For the binary model, predictions were then binarised, with each value above 0.5 becoming 1.0, and each value below 0.5 becoming 0.0 (values at exactly 0.5 were treated as missed trials for the binary model). Finally, the outcomes of each trial were adjusted to be in ‘contingency space’: i.e., any trial where cue 1 was paired with a resistance and cue 2 with no resistance was coded as 1, and any trial where the cue-outcome pairing was reversed was coded as 0, as previously described (Harrison et al., 2021; Iglesias et al., 2013). Notably, contingency space coding does not depend on which binary value is assigned to coupled cue-outcome pairs and is equivalent to the case of running two models in parallel, one for each outcome (for details, see supplementary material to Iglesias et al. (2013)). This type of coding was possible because of the fixed coupling of contingencies in our task (see Section 3.2.2 for more detail), which the participants were made explicitly aware of. However, this method does assume that learning is the same for resistance and no resistance trials. To address this assumption, we extended our model to separate the learning parameters for resistance and no resistance trials (see Section 3.5.2).

### 3.4 Task validation

In order to validate that participants were completing the task as expected, the proportion of correct (binarised) responses on each trial across all participants was compared with data from Harrison et al. (2021), which investigated a larger cohort of participants using the binary prediction version of the BLT. This allowed us to verify that overall, participants understood the task and behaved in line with previous findings from a larger cohort.

### 3.5 Associative learning model

The Rescorla-Wagner associative learning rule was utilised (Rescorla et al., 1972), where the predicted outcome for a given trial *v*_*t*+1_ is based on the predicted outcome for the previous trial *v*_*t*_ as well as the prediction error for the previous trial *δ*_*t*_ scaled by the learning rate of the participant *α*:

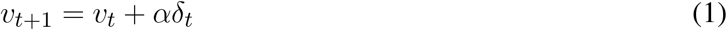

where the prediction error is the difference between the actual outcome *o*_*t*_ and the predicted outcome *v*_*t*_for the previous trial:

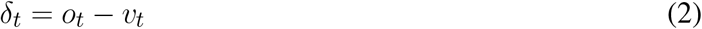

Previous research has employed similar models on binary BLT data (see Harrison et al. (2021)), where participants had two choices (Resistance/No Resistance) when asked to predict the outcome of each trial. However, in the current version of the BLT, participants were asked to predict the outcome of each trial on a sliding scale with values from 0.0 (Definitely No Resistance) to 1.0 (Definitely Resistance). These continuous data therefore includes a direct measure of the participant’s certainty in their predictions, rather than requiring this to be inferred from fitted model trajectories. In order to apply the model to the participants’ observed responses, the Rescorla-Wagner learning model (Eq. 1) was paired with two different observation models - one using binarised responses (for comparisons with previous versions of the task) and the other continuous responses (henceforth referred to as the binary model and the continuous model respectively). In either case, the participant’s responses (i.e. binary or continuous predictions about trial outcome) and outcomes (binary: absence or presence of resistance) from each trial were modelled by estimating a subject-specific learning rate (*α*) for the participant (see Eq. 1 above). The specifics of each observation model are discussed in Section 3.5.1 below.

#### 3.5.1 Observation models

The observation model translates the predicted outcome, as provided by the learning model, into predicted behaviour (i.e. moving the slider to a certain position for the next trial in the case of continuous data, or a binary decision for the binarised data).

##### 3.5.1.1 Binary model

The binary model uses a softmax function as the observation model, which translates the estimates obtained by the learning model into the probability of choosing a given action - in this case, deciding whether a given stimulus predicts resistance or no resistance. The softmax function used by the binary model can be represented as follows:

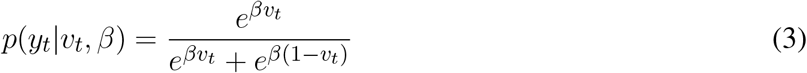

Where *p*(*y*_*t*_|*v*_*t*_, *β*) represents the estimated probability of choosing a given binary prediction response *y*_*t*_ given the predicted outcome *v*_*t*_, *β* is a static parameter that determines the steepness of the gradient of the softmax function, and *v*_*t*_ is the value calculated by the learning model. The parameter *β* can be altered to represent more or less noise in the decision-making process, with a higher beta resulting in a steeper softmax gradient, and thus more deterministic behaviour.

##### 3.5.1.2 Continuous model

As the continuous model allows for predictions to occur on a continuous scale rather than having to make a binary decision, the value obtained from the learning model for a given trial *v*_*t*_ is represented by the observed continuous prediction response *y*_*t*_. The likelihood of the data is derived using a beta distribution, given that the responses lie in the range [0,1], and that the beta distribution is an adequate model for continuous bounded data, such as proportions or probabilities. Here, we use a formulation which re-parameterises the usual shape parameters of the beta distribution in terms of parameters for mean and dispersion (Paolino, 2001). Moreover, we make dispersion a group parameter, *ϕ*, where higher *ϕ* represents less noise.

Specifically, this is implemented by re-parameterising the shape parameters of the standard beta distribution (specified by *a* and *b*, both vectors of length *n*_*trials*_ *· n*_*subjects*_) in terms of mean *μ* (a vector of length *n*_*trials*_ · *n*_*subjects*_, which contains the values obtained from the learning model (see equation 1) for each subject and trial) and a scalar dispersion parameter *ϕ* (which is constant across subjects and trials), as follows (Paolino, 2001):

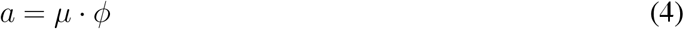

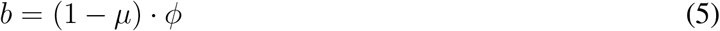

As a result, large values of *ϕ* result in a tighter distribution while smaller values result in a wider distribution. In other words, less consistent (i.e., noisier) responses across subjects are reflected by a smaller value of *ϕ*. Here we estimate a single value of *ϕ* across all subjects, though we note that subject-specific values could in principle be estimated to allow for differences in response consistency across subjects.

#### 3.5.2 Dual learning rate model

To investigate whether there were learning differences between positive and negative valence stimuli, a variation of the model was created that used an altered version of the Rescorla-Wagner algorithm. This version is equivalent to equations 1 and 2 above, except that it contains two learning rates: *α*_*p*_ and *α*_*n*_. Which learning rate is used for the update on a given trial depends on the stimulus type *s* that occurred during that trial - no breathing resistance is considered a positive valence stimulus (represented as *s*_*t*_ = 0) while breathing resistance is considered a negative valence stimulus (represented as *s*_*t*_ = 1). The state equation for the dual learning rate model is thus as follows:

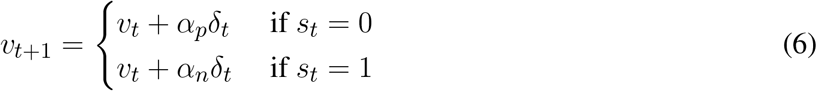

where the predicted error *δ*_*t*_ is calculated as in equation 2.

### 3.6 Model testing

#### 3.6.1 Parameter estimation and prior selection

Both models used maximum a posteriori (MAP) methods to obtain parameter estimates (using single-start optimisation and the L-BFGS-B algorithm (Byrd et al., 1995; Zhu et al., 1997)), with the following priors: for *α* (as well as for *α*_*p*_ and *α*_*n*_ for the dual learning rate model) the prior mean was 0.34 and variance was 0.88 and for *β* the prior mean was 4.21 and variance was 1.75 (all specified in native space using a Gaussian distribution, with *α* values bounded between 0.0 and 1.0). These prior distributions were calculated using the binary model to obtain Maximum Likelihood Estimates (MLE) from data of a separate group of eight pilot participants, originally used for the Harrison et al. (2021) study. The initial value of *v* (the predicted outcome) was fixed at 0.5 (representing complete uncertainty of the outcome) for both models. The code for used for model inversions can be found at https://github.com/IMAGEotago/Katja-BLT-code.

#### 3.6.2 Simulation and parameter recovery

To test and validate the models, we first simulated responses to the BLT for hypothetical participants with a range of different learning rates. This was performed 500 times for each version of the model (as pre-specified in the analysis plan), using a randomly generated learning rate each time - drawn from a truncated normal distribution with a mean of 0.34, a variance of 0.88, with lower and upper bounds of 0.0 and 1.0 respectively. Each simulation was performed using the same sequence of outcomes for each trial as in the BLT. The simulations were repeated for four different *β* values (of the softmax response function) for the binary model. For the continuous model, noise drawn from a Gaussian distribution with a mean of 0.0 and a standard deviation *σ* was added to the simulated prediction responses in order to reflect the noise inherent in real-world behavioural data. Once noise was added, the observed values were constrained such that the simulated prediction responses *y*_*t*_ remain within 0.0 and 1.0 by bounding upper and lower values. Similarly to the binary model, simulations for the continuous model were repeated for four different *σ* values (representing different levels of noise). An example of the simulated trajectories can be seen in Figure 3.

**Figure 3.**
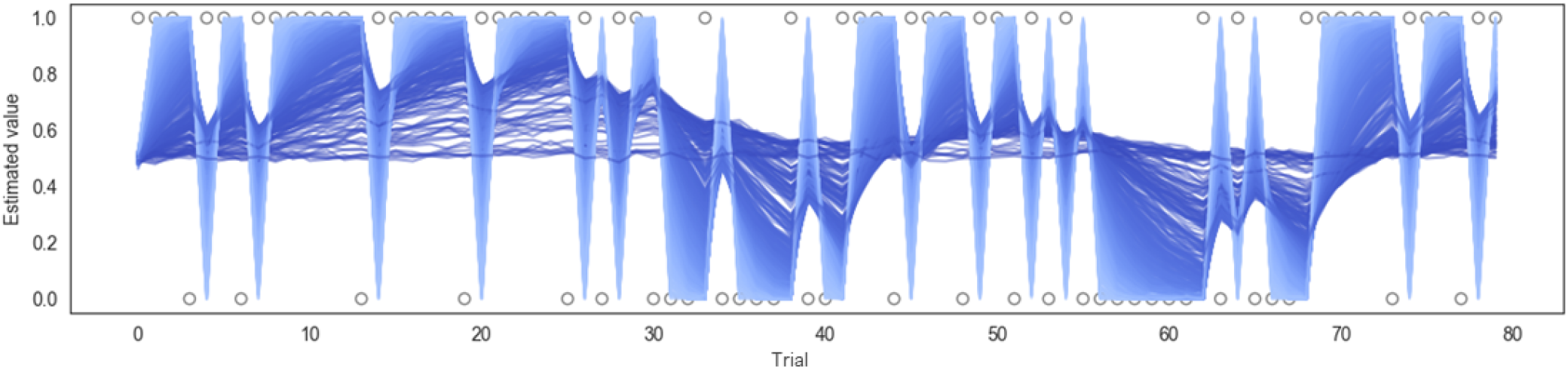
Example of simulation results using the continuous model for 500 subjects (*σ* = 0.01). Each line represents a different simulated subject, with a lighter blue line indicating a high *α* and a darker blue line indicating a lower *α*.

The data generated from each simulation was then fitted to assess how accurately the model parameters could be recovered (Wilson and Collins, 2019). 10 runs of simulation and parameter recovery were completed at each noise level (simulating 500 subjects each time), to ensure consistency was high across simulation runs. The results for parameter recovery are presented in Section 4.2.1 below.

#### 3.6.3 Model inversions on empirical data

Following successful parameter recovery using synthetic data for both models, the empirical data gathered from the BLT for each participant were used to invert each model. The model input consisted of the predictions made by participants on each trial along with the outcomes of each trial (resistance or no resistance, with both predictions and outcomes in contingency space; see Section 3.3). From this input, the models estimate a learning rate (*α*) for each participant, as well as either a *β* value (of the softmax response function) when using the binary model or a group-level dispersion parameter *ϕ* when using the continuous model. The same process was used to obtain estimates for *α*_*p*_ and *α*_*n*_ values (and *β* for the binary version) for the dual learning rate model. The results for each model for all participants are presented in Section 4.2.2 below.

#### 3.6.4 Model validation

The next step for validating the models was to verify that they provided useful information from fitting the participants’ data. In order to do this, both the binary and continuous model fits to participant data were compared to a null version of the respective model. This null model works the same as the binary or continuous model except that the learning rate *α* was fixed at 0.0. It therefore represents a condition where no learning occurs (i.e. *v*_*t*_ is clamped to 0.5) and all input is attributed to noise. The binary and continuous model were validated by comparing the model fits to the respective null model fits using the Bayesian Information Criterion (BIC; Schwarz, 1978) and the Akaike Information Criterion (AIC; Akaike, 1973). Both the BIC and the AIC are ways of determining which model is ‘best’ in terms of the trade-off between goodness of fit and model simplicity (Kuha, 2004). Although this inverts the original definition of BIC (Schwarz 1978), one way to include the definitions of both BIC and AIC in one expression is:

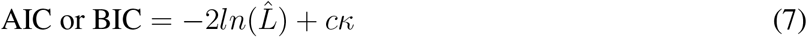

Here 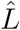 is the log likelihood of the data given the parameter estimates, *κ* is the total number of parameters in the model (as a proxy of model complexity), and *c* is a penalty coefficient. For the BIC, *c* = *ln*(*n*) where *n* is the number of observations, while for the AIC, *c* = 2 (Vrieze, 2012). Given the above formulation (Equation 7), the model with the smaller BIC or AIC value is considered to be the better model in terms of how well it minimises information loss. A conversion into Bayes factors can be used to quantify the degree to which one model is preferred (Penny et al., 2004; Wagenmakers, 2007). Since BIC and AIC have different notions of model complexity, we used both of them to compare the binary and continuous models against the respective null models. It should be noted that model comparison by BIC and AIC is only valid when comparing models fit to the same data, therefore they cannot be used to compare directly between the binary and continuous models.

In addition to this, a second validation was performed by analysing how well model fits correlated to actual participant behaviour at the group level. To do this, the average prediction trajectories fitted by the model were compared to the actual predictions of the participants (averaged across participants for each trial) using a Pearson correlation to determine how well they were aligned. Results from both of these tests are presented in Section 4.2.3 below.

#### 3.6.5 Comparisons between single and dual learning rate models

To determine whether the dual learning rate models were capturing useful additional information to the single learning rate models, the BIC and AIC were used to compare the binary and continuous versions of the two models (using the same approach as described in Section 3.6.4 above). Results are presented in Section 4.2.4 below.

#### 3.6.6 Exploratory correlations with questionnaires

Finally, we investigated how the information provided by these models could be used to explore how interoceptive learning may relate to both measures of mental health and subjective interoception. To do this, exploratory non-parametric Spearman rank correlations were performed between the questionnaire scores (STAI-S, STAI-T, GAD-7, ASI-3, CESD, PANAS-P/N, FSS, CD RISC-25, GSE, MAIA, PCS-B,

PVAQ-B), the learning rates estimated by the different models, and the average prediction certainty of each participant using their continuous prediction data. Due to the exploratory nature of these correlations, findings are uncorrected for multiple comparisons. The findings are reported in Section 4.3 below.

## 4 RESULTS

### 4.1 Task validation

To investigate the consistency in task performance in the current study with the previous binary version of the BLT, the proportion of correct responses on each trial across all participants for the current study was firstly compared to the larger cohort of previous data (Harrison et al., 2021) (see Figure 4). A strong correlation was found between the proportion of correct responses at each trial across the two cohorts, with a Pearson’s correlation coefficient of *r* = 0.85 (*p* = 5.18*e*^*−*^23). This indicates that overall, participants responded to the new version of the task in a similar manner to the larger cohort of participants used in the previous study.

**Figure 4.**
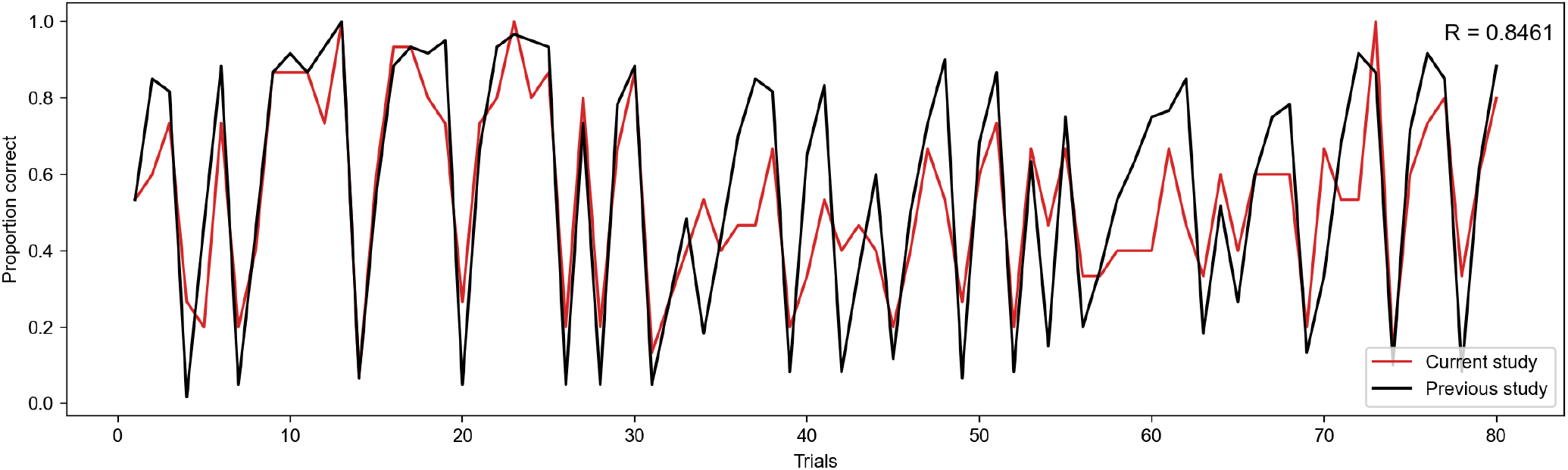
Proportion of correct responses for each trial across the current study cohort (red line) and the previous study cohort (black line).

### 4.2 Model results

#### 4.2.1 Simulation and parameter recovery

##### 4.2.1.1 Binary model

For the binary model, simulation and parameter recovery was performed at four different values of *β* (steepness of the gradient of the softmax function, representing decision noise): 1, 2, 4, and 8, with 10 simulation runs for each value of *β* and 500 simulation subjects in each run. The correlation between the simulated values of *α* and the recovered values of *α* was averaged across the 10 runs for each different value of *β* by converting the r-values to z-values, taking the mean, and then converting back to r-values. Figure 5 shows the results for a representative run for each different value of *β*.

**Figure 5.**
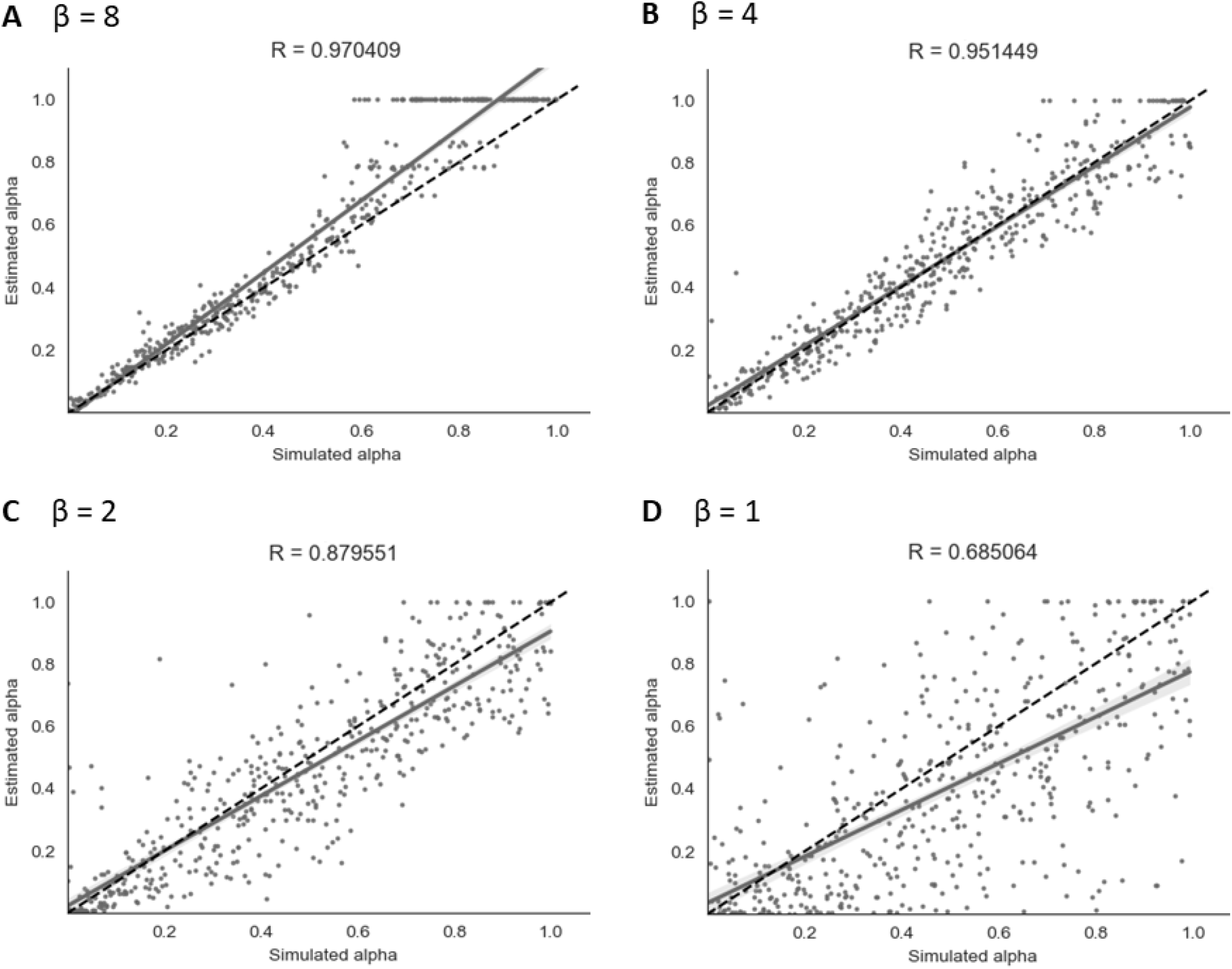
Parameter recovery of *α* for the binary model at four different values of *β*. Data points shown were taken from one representative simulation run.

As shown in Figure 5, parameter recovery was very successful for higher values of *β*, with an average correlation between estimated and simulated *α* of *r* = 0.97 for *β* = 8, *r* = 0.95 for *β* = 4 and *r* = 0.88 for *β* = 2. There was still a moderate correlation between estimated and simulated *α* of *r* = 0.68 for *β* = 1. As can be seen in Figure 5, there appears to be a ceiling effect for recovered *α* values above *α*≈ 0.6 when there is less noise present. This finding has been reported previously with binary data (Harrison et al., 2021). The simulated *β* values were also successfully recovered, with the exception of *β* = 8 where the recovered *β* value was lower than the simulated *β* value. A visualisation of this recovery can be found in Section 2 of the Supplementary Materials.

##### 4.2.1.2 Continuous model

Similarly to the binary model, simulation and parameter recovery was performed at four different values of *σ* (the standard deviation of the added noise) - 0.05, 0.1, 0.2, and 0.4, with 10 simulations runs and 500 simulated subjects for each value of *σ*. Mean r-scores for each level of noise were calculated as for the binary model above. Examples of results from a representative run for each level of noise are shown in Figure 6.

**Figure 6.**
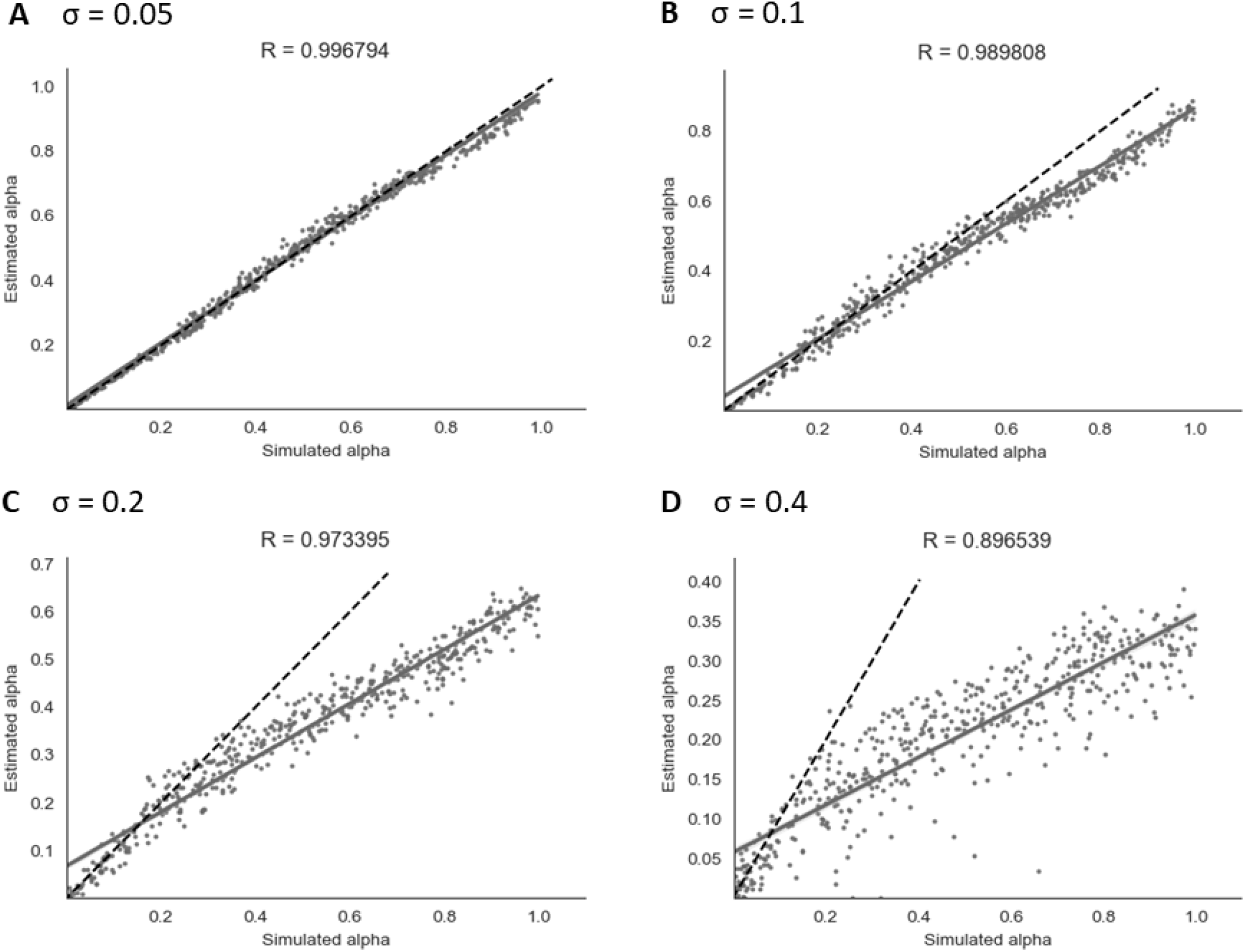
Parameter recovery of *α* for the continuous model at four different values of *σ*. Data points shown were taken from one representative simulation run.

As shown in Figure 6, parameter recovery showed a significant correlation between simulated and recovered *α* values at all levels of noise. The average correlation for each noise level was *r* = 1.00 at *σ* = 0.05, *r* = 0.99 at *σ* = 0.1, *r* = 0.97 at *σ* = 0.2, and *r* = 0.90 at *σ* = 0.4. At higher noise values, an under-estimating bias became apparent, with recovered values being estimated in a lower range than simulated values. Increasing the Gaussian noise (via increased *σ*) in simulations produced the expected reductions in the recovered group-level dispersion parameter *ϕ*, and the values used for simulation can be found in Section 2 of the Supplementary Materials.

##### 4.2.1.3 Dual learning rate model

Simulation and parameter recovery was also carried out for both the binary and continuous versions of the dual learning rate model, using the same method as explained above. Parameter recovery showed similar results as the single learning rate model for the continuous version, with strong correlations between simulated and recovered *α* values at all levels of noise: at *σ* = 0.4 (*r*(*α*_*p*_) = 0.84, *r*(*α*_*n*_) = 0.88), *σ* = 0.2 (*r*(*α*_*p*_) = 0.94, *r*(*α*_*n*_) = 0.96), *σ* = 0.1 (*r*(*α*_*p*_) = 0.98, *r*(*α*_*n*_) = 0.99), and *σ* = 0.05 (*r*(*α*_*p*_) = 0.99, *r*(*α*_*n*_) = 1.00). Similar results were obtained with the binary version - see Supplementary Material Section 4.1 for further detail and a graphical representation of the parameter recovery results for the dual learning rate model.

#### 4.2.2 Model inversions on empirical data

##### 4.2.2.1 Binary and continuous models

Both the binary and continuous models were used to estimate a learning rate (*α*) for each of the participants (as well as a *β* value for the binary model or a *σ* value for the continuous model) and produce a corresponding prediction trajectory. The prediction trajectories for each of the fitted learning rates from both models are displayed in Figure 7. Figure S4 in the Supplementary Material shows a comparison of individual response data to model-fitted trajectories for a range of learning rates.

**Figure 7.**
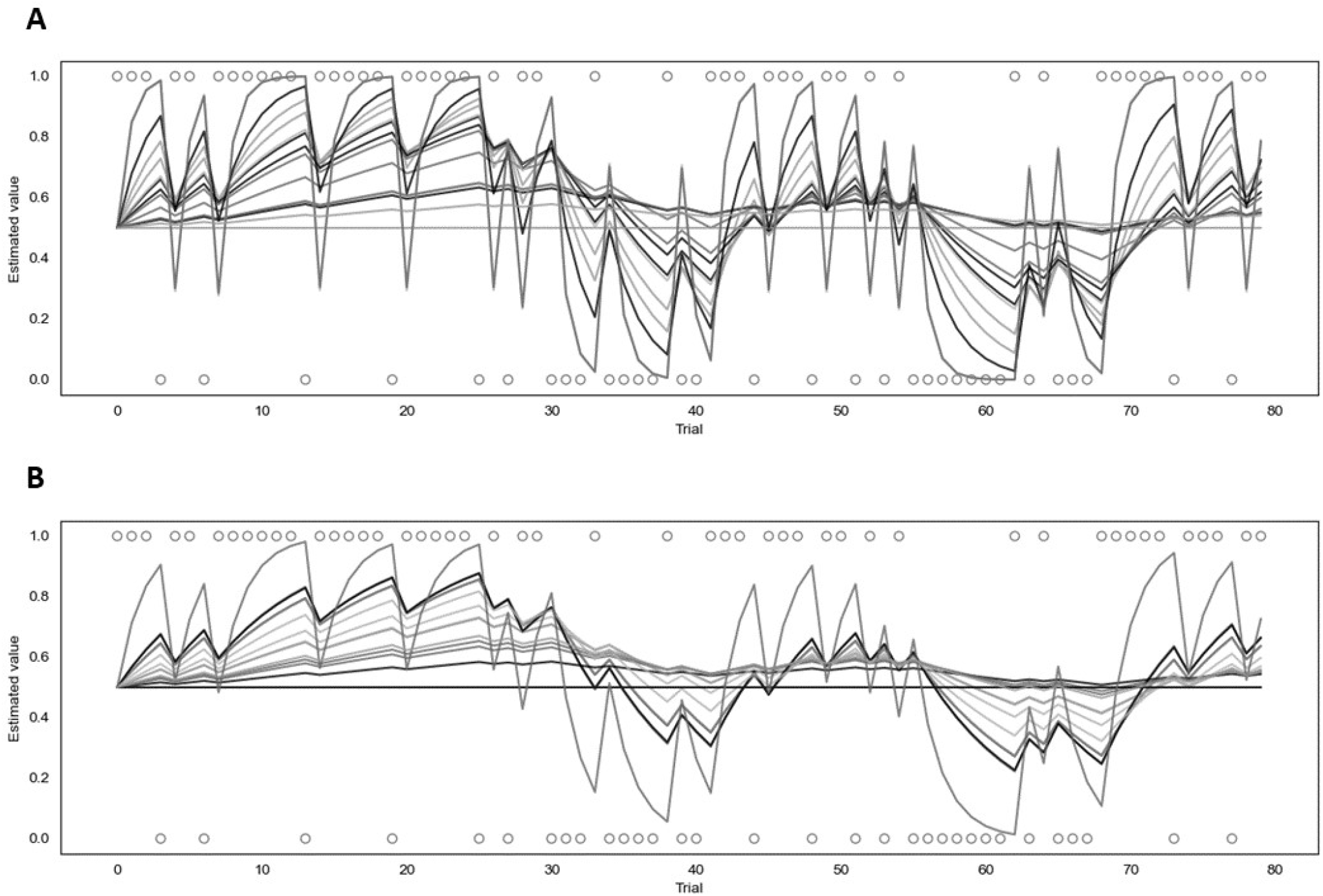
Fitted prediction trajectories for each participant. **(A)** represents results from the binary model, **(B)** represents results from the continuous model. Circles represent trial outcomes.

As can be seen in Figure 7, the *α* values fitted by each of the models varied across participants. For the continuous model, the mean *α* value was 0.08 with a standard deviation of 0.10 and the value of the group parameter *ϕ* was 0.04. For the binary model, the mean *α* value was 0.18 with a standard deviation of 0.22, and the mean *β* value was 4.68 with a standard deviation of 2.00. Learning rates fitted by the binary and continuous model were compared using a t-test, which indicated that the learning rates fitted by the continuous model were significantly lower than those fitted by the binary model (*p* = 0.03).

##### 4.2.2.2 Dual learning rate model

For the dual learning rate model, both the binary and continuous versions were used to estimate two learning rates (*α*_*p*_ and *α*_*n*_) for each of the participants. The corresponding prediction trajectories generated for each participant are displayed in Figure S7 in the Supplementary Material. For the continuous version, the mean value of *α*_*p*_ was 0.09 (standard deviation of 0.11), the mean value of *α*_*n*_ was 0.07 (standard deviation of 0.11), and the group parameter *ϕ* was 0.05. For the binary version, the mean value of *α*_*p*_ was 0.25 (standard deviation of 0.26), the mean value of *α*_*n*_ was 0.19 (standard deviation of 0.26), and the mean value of *β* was 4.31 (standard deviation of 2.49).

#### 4.2.3 Comparison against null models and validation

##### 4.2.3.1 Binary and continuous models

To test whether the binary and continuous models were providing useful information when fitting participants’ data, they were each compared to their respective null models (see Section 3.6.4 for a full explanation). Both models achieved a smaller BIC and AIC score (indicating a better fit) than their null model counterparts. The binary model produced a BIC of 1253 and an AIC of 1252 compared to the binary null model which produced a BIC and AIC of 1691. Calculations of the Bayes factor indicate the binary model is preferred over the null model with a Bayes factor of 1.36 × 10^95^. The continuous model produced a BIC of -1315 and an AIC of -1316 compared to the continuous null model which produced a BIC and AIC of -1070, leading to the continuous model being preferred with a Bayes factor of 1.57 ×10^53^. Both of these Bayes factors indicate very strong evidence favoring the binary and continuous models (Raftery, 1995).

For the group level validation, an average predicted trajectory was created by calculating the mean predictions generated by the model across all participants for each trial. This was then compared to the mean of the observed prediction values entered by each participant on each trial, using a Pearson correlation. This procedure was performed for both the binary and continuous model, with the binary model being compared to the mean of the binarised observed prediction values that were used as the input for that model.

The results are presented in Figure 8. The mean observed predictions and mean modelled predictions showed a strong correlation for both the binary (*r* = 0.80, *p* = 8.18*e*^*−*19^) and continuous (*r* = 0.79, *p* = 1.41*e*^−18^) models.

**Figure 8.**
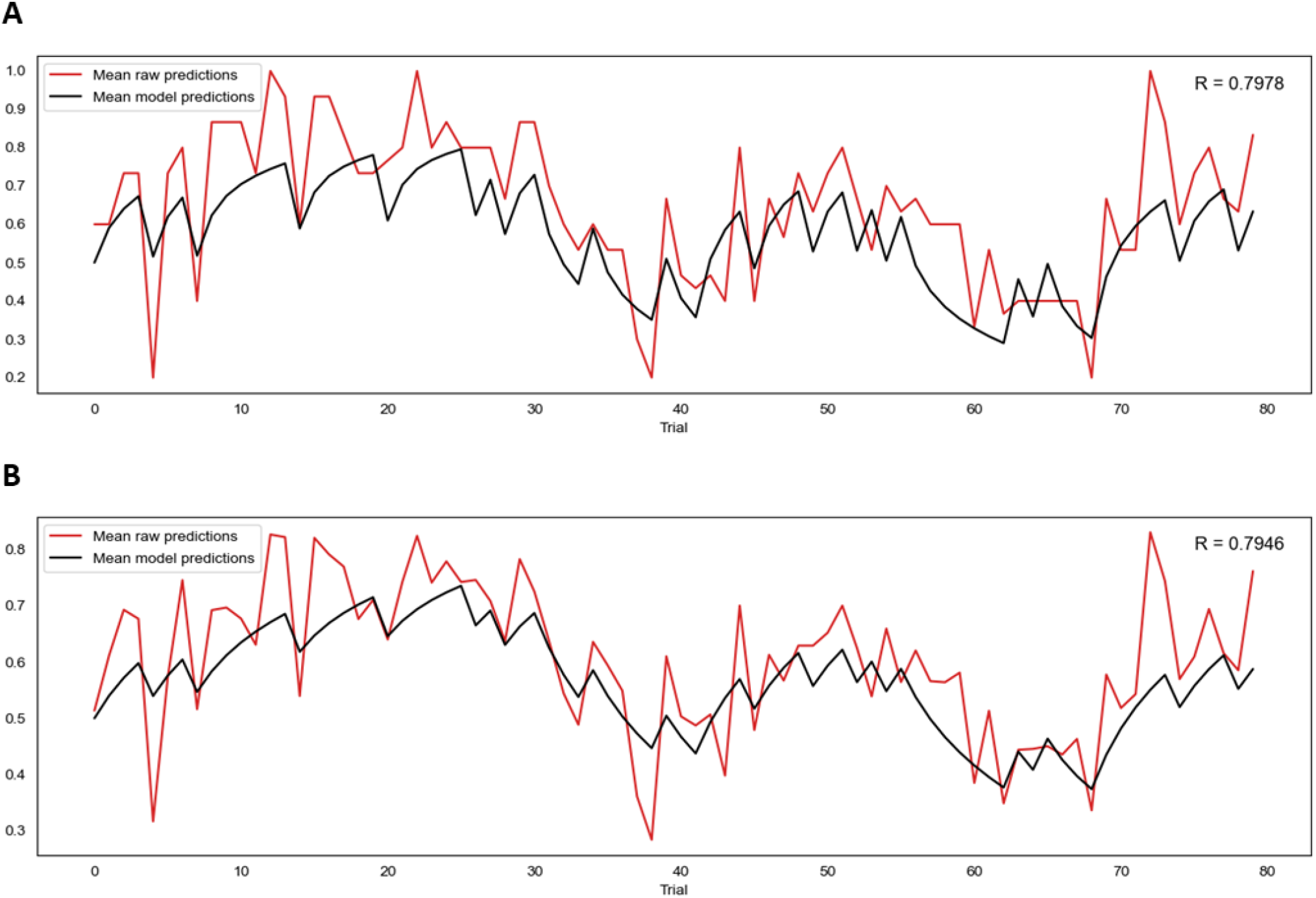
**(A)** Correlation between mean model predictions across all participants and mean observed prediction values for the binary model. **(B)** Correlation between mean model predictions across all participants and mean observed prediction values for the continuous model.

#### 4.2.4 Comparisons between single and dual learning rate models

To determine whether the dual learning rate binary and continuous models were explaining the data better than their single learning rate counterparts, the models were compared using the BIC and AIC metrics. For the continuous models, the dual learning rate version produced a BIC of -1298 and an AIC of -1299 when run on the participants’ data compared to a BIC of -1315 and an AIC of -1316 for the single learning rate version, resulting in a Bayes factor of 6311 in favor of the single learning rate version. Results were similar for the binary models, with the dual learning rate version producing a BIC of 1262 and an AIC of 1259, while the single learning rate version had a BIC of 1253 and an AIC of 1252, resulting in a Bayes factor of 70.1 in favor of the single learning rate version. These comparisons provide strong to very strong evidence that the both the binary and continuous single learning rate models represent a better trade-off between accuracy and complexity than the corresponding dual learning rate models.

### 4.3 Questionnaire results

Participants completed a number of questionnaires measuring affective and interoceptive qualities (see Section 3.2.1 for full list). Median scores and interquartile ranges (IQR) across participants were calculated for each questionnaire and are presented in Table 1.

**Table 1.** Questionnaire scores.

|  | Median (IQR) |
| --- | --- |
| STAI-T | 36.5 (18.75) |
| STAI-S | 33 (18.75) |
| GAD-7 | 2.5 (4.75) |
| ASI-3 | 17.5 (24.75) |
| CESD | 10 (7.75) |
| PANAS-P | 33.5 (8.5) |
| PANAS-N | 18 (10.25) |
| FSS | 35 (17.75) |
| CD-RISC 25 | 71 (18.25) |
| GSE | 31 (19) |
| MAIA | 21.29 (6.37) |
| PCS-B | 11.5 (29) |
| PVAQ-B | 32.5 (8.5) |

Exploratory correlations were then performed between the questionnaire scores and the estimated learning rates obtained from the continuous (*α* (c)) and binary (*α* (b)) models, as well as the average certainty across trials for each participant from the continuous prediction data. The results are presented in a correlation matrix in Figure 9. Results are uncorrected as the correlations are exploratory in nature.

**Figure 9.**
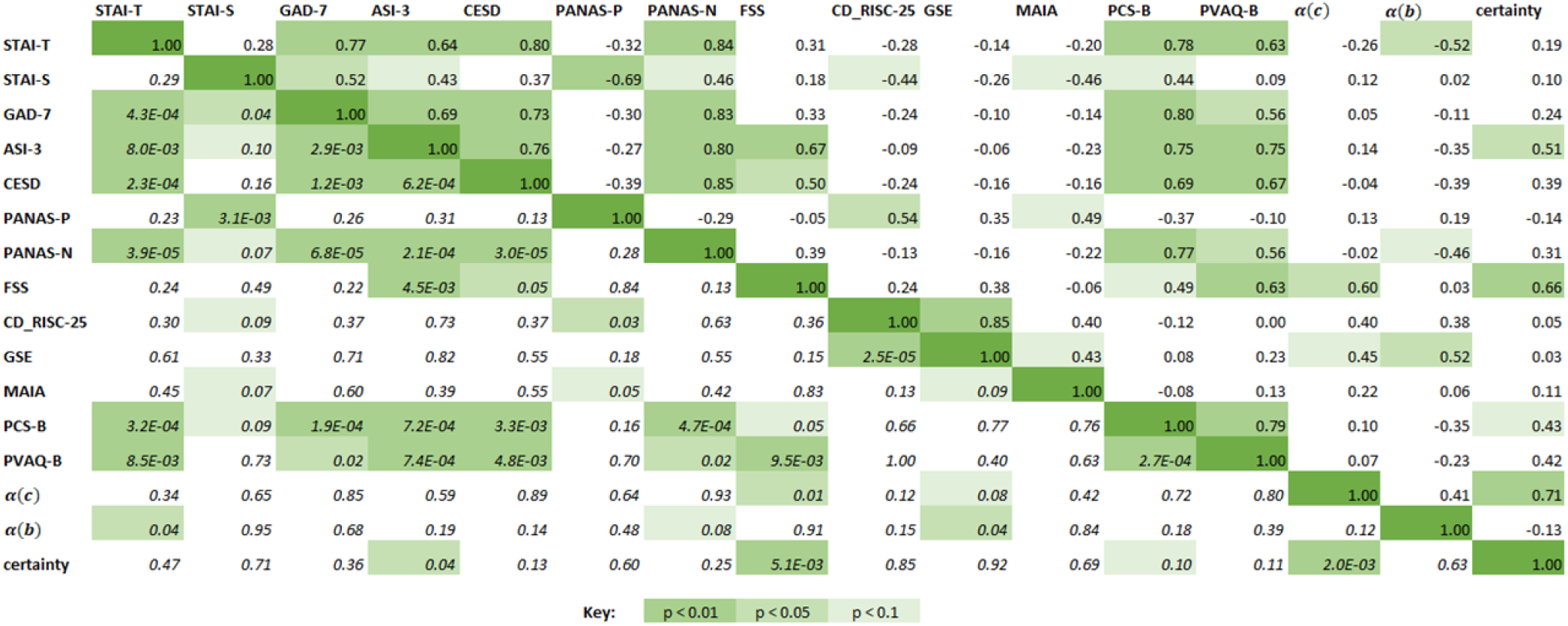
Correlation matrix for questionnaire scores, learning rates from the continuous and binary models, and average certainty. The upper right half of the matrix contains the Spearman correlation coefficients, while the italicised fields represent the corresponding *p*-values for each correlation score. Green highlighted fields indicate significant results at *p <* 0.1, *p <* 0.05, *p <* 0.01 with a darker green indicating a more significant result.

As can be seen in Figure 9, significant correlations were observed between the questionnaire measures of anxiety, depression and affect (STAI-T, STAI-S, GAD-7, ASI-3, CESD, PANAS-P and PANAS-N), and some of these were additionally related to measures of interoceptive qualities (MAIA, PCS-B, and PVAQ-B) as expected. Fatigue severity (FSS) was positively correlated with anxiety sensitivity (ASI-3, *r*_*s*_ = 0.67, *p* = 0.005), depression symptoms (CESD, *r*_*s*_ = 0.50, *p* = 0.05), and increased vigilance around breathlessness (PVAQ-B, *r*_*s*_ = 0.63, *p* = 0.01). Between the BLT data and questionnaire scores, there was a moderate correlation between FSS scores and both average certainty values (*r*_*s*_ = 0.66, *p* = 0.01) as well as continuous model learning rates (*r*_*s*_ = 0.60, *p* = 0.01). There was also a significant correlation between average certainty values and ASI-3 scores (*r*_*s*_ = 0.51, *p* = 0.04). Binary model learning rates were negatively correlated with STAI-T (*r*_*s*_ = −0.52, *p* = 0.04) and positively correlated with GSE scores (*r*_*s*_ = 0.52, *p* = 0.04).

## 5 DISCUSSION

This work has provided an advancement on the interoceptive learning paradigm (BLT) used in Harrison et al. (2021) by incorporating continuous response data, thus providing a more direct measure of prediction certainty. Additionally, an extension of the computational model tested whether stimuli valence had an effect on learning rate, investigating whether separate learning rates for positive and negative stimuli should be considered when fitting behavioural data. Data from a novel continuous version of the BLT was gathered from a cohort of 16 healthy participants, and behavioural data from this cohort were compared to that gathered by Harrison et al. (2021). Importantly, a close correlation was observed in participant performance (i.e. percentage of correct predictions) between the two cohorts, indicating that participants were similar in their overall task performance when moving from binary to continuous predictions.

When considering model validity using simulated data, both the binary and continuous models provided significant parameter recovery across varying levels of noise. It should be noted that for the continuous model, constrained Gaussian noise was added to the simulated trajectories. This noise added variability to simulated responses, but was constrained such that prediction values remained between 0 and 1. However, these Gaussian noise assumptions were not implemented during model inversion (see Section 3.5.1), with variability instead captured by a group-level dispersion parameter, *ϕ*.

For both binary and continuous data, comparisons to the respective null models using the BIC and AIC indicated that our trial-wise learning models were greatly superior, indicating that they were capturing useful information from participants’ responses. Furthermore, the average modeled prediction trajectories from both models were highly correlated with the average observed predictions, suggesting that both models did well at representing the behaviour of participants at a group level.

No significant correlation was observed when comparing the estimated learning rate parameters fit by each of the models to the empirical data. Furthermore, the continuous model generally produced smaller *α* values in comparison to the binary model. This may be influenced by both the under-estimation bias and the reduced variability in the estimated learning rates observed in the continuous vs. binary model simulation results. In addition, two of the participants had estimated learning rates below 0.0001 by one or both of the models, which would indicate that little to no learning occurred during the task. It is therefore possible that these participants did not properly understand the task or failed to follow instructions, which future studies using this task may need to further accommodate for.

In addition to incorporating continuous predictions, we also explicitly tested the model assumption that a single learning rate was able to adequately capture participant behaviour across the resistance and no-resistance stimuli. One potential issue with a single learning rate model is that it assumes that participants learn at the same rate from negative (i.e. resistance) as for positive (i.e. no resistance) outcomes, as participants were explicitly told that cues act as a pair. However, previous research has indicated that people may learn differently from negative compared to positive stimuli, and this may be influenced by factors such as anxiety (Aylward et al., 2019; Khdour et al., 2016). To test whether learning differed according to stimuli valence in the BLT, the model was extended to introduce a dual learning rate algorithm that estimated separate learning rates following resistance and no-resistance trials. Overall, both binary and continuous models that incorporated a dual learning rate produced results consistent with their single learning rate counterparts. However, the results from comparing the BIC and AIC for each of the models suggest that the single learning rate models are preferred over the dual learning rate models in terms of accuracy-complexity trade-off. Therefore, as the dual learning rate model did not convey an advantage to explaining participants’ behaviour, a meaningful difference in learning between positive and negative stimuli for this task is unlikely. Thus, the original assumptions made for the single learning rate model are likely adequate, although a larger sample size would allow for further validation of this result.

Finally, one further advantage of the continuous version of the BLT is the additional information captured in the observed response data. Binary data records only the direction of the prediction (i.e. resistance or no resistance) while continuous responses additionally contain information about the degree of certainty in the prediction. This information could only be inferred from the binary data through a model, while the continuous data provides a more direct readout from participants, allowing prediction certainty to be measured independently of any model assumptions. We investigated whether learning rate parameters fit by either model and/or the observed prediction certainty from the continuous task were related to questionnaire measures of both affective and interoceptive qualities. Fatigue severity (as measured by the FSS) was found to be positively correlated with both the learning rate fit by the continuous model and with average response certainty, but not the learning rate from the binary model. However, learning rates from the binary model were correlated with questionnaire measures of trait anxiety and self-efficacy. Additionally, there was a significant correlation between average certainty and anxiety sensitivity (as measured by the ASI-3). These results indicate that the continuous data provide us with different information compared to when we only consider binary decisions. However, the binary information is not lost in this version of the task, allowing the data to be analysed in multiple ways. Overall, these preliminary findings demonstrate that valuable information can be gained from including a direct measure of response certainty within interoceptive learning tasks when investigating the relationship between interoception and mental health.

## 6 CONCLUSION

This report has presented a method of incorporating continuous response data into the interoceptive learning paradigm (BLT) introduced by Harrison et al. (2021), and presented a suitably extended response model for a Rescorla-Wagner learning model of the measured data. Furthermore, it tested whether assuming a single learning rate, regardless of stimulus valence, was adequate for the BLT, or whether an extended model with two separate learning rates would be advantageous. Both binarised and continuous data from a pilot cohort who completed the modified BLT were fit using a Rescorla-Wagner learning model. Both models performed well on simulations (recovery analyses) and fitting the empirical data. While there was some variability of individual model fits, the continuous model accurately captured behaviour at the group level. Our preliminary analysis indicates that this new model for continuous response data during the BLT could be useful for investigating the relationship between mental health and interoceptive learning, with the additional information regarding prediction certainty demonstrating significant relationships with fatigue and anxiety sensitivity scores. The task and model described here provide a useful toolbox for future investigations into interoceptive learning (Frässle et al., 2021).

## CONFLICT OF INTEREST STATEMENT

The authors declare that the research was conducted in the absence of any commercial or financial relationships that could be construed as a potential conflict of interest.

## AUTHOR CONTRIBUTIONS

OKH and BRR designed the study, using methodology developed by OKH, KES and AJH. Data collected for this study was completed by KB and OKH. Model implementation and extension was performed by KB, with input from OKH, TW and AJH. KB and OKH wrote the manuscript with edits from all remaining authors.

## FUNDING

This study was supported by a Rutherford Discovery Fellowship awarded by the Royal Society of New Zealand to Dr Harrison, with additional support provided by the School of Pharmacy and Department of Psychology at the University of Otago. Ms Brand was supported by a University of Otago Master’s scholarship, and the writing of the manuscript was supported by the University of Otago Postgraduate Publishing Bursary (Master’s). Dr Stephan acknowledges support from the René and Susanne Braginsky Foundation and the ETH Foundation.

## ACKNOWLEDGMENTS

The authors would like to thank Sophie Toohey for her assistance with participant recruitment and data collection as part of a wider study. Thanks also to our clinical Pharmacists Emma Smith, Lauren Smith and Tara Wheeler for their support.

## DATA AVAILABILITY STATEMENT

The datasets generated for this study can be made available on request.

## FIGURE CAPTIONS

**Figure 1**. Schematic of the inspiratory resistance circuit used. Participants were connected to the circuit via a facemask covering their mouth and nose which was attached to a bacterial and viral filter (labelled C). A sampling line to a pressure transducer (labelled O) and amplifier was used to record inspiratory pressure, and a spirometer (labelled E) was used to measure inspiratory flow. Inspiratory resistances were induced automatically via the stimulus computer, controller box and solenoid valve (labelled L) to redirect unobstructed air from the environment to the PowerBreathe device (labelled K), which provided a set magnitude of resistance. Figure adapted from Rieger et al. (2020).

**Figure 2**. (A) An overview of the structure of a single trial in the version of the BLT used in Harrison et al. (2021). (B) An overview of the structure of a single trial in the version of the BLT used for the current study. The main changes are that the cue/prediction phase was changed to allow continuous predictions via a slider (with a longer timeframe for this phase to compensate), while the report phase was change to a binary yes/no response to simplify this decision for participants. Additionally the pause before the stimulus was removed after lengthening the cue/prediction phase. Figure adapted from Harrison et al. (2021).

**Figure 3**. Example of simulation results using the continuous model for 500 subjects (*α* = 0.01). Each line represents a different simulated subject, with a lighter blue line indicating a high *α* and a darker blue line indicating a lower *α*.

**Figure 4**. Proportion of correct responses for each trial across the current study cohort (red line) and the previous study cohort (black line).

**Figure 5**. Parameter recovery of *α* for the binary model at four different values of *β*.

**Figure 6**. Parameter recovery of *α* for the continuous model at four different values of *σ*.

**Figure 7**. Fitted prediction trajectories for each participant. (A) represents results from the binary model, (B) represents results from the continuous model. Circles represent trial outcomes.

**Figure 8**. Correlation between the *α* values fitted by the continuous and binary models for each participant. Light grey dots signify participants whose binary model fits did not pass the LRT.

**Figure 9**. (A) Correlation between mean model predictions across all participants and mean raw prediction values for the continuous model. (B) Correlation between mean model predictions across all participants and mean raw prediction values for the continuous dual learning rate model.

**Figure 10**. Correlation matrix for questionnaire scores, learning rates from the continuous and binary models, and average certainty. The upper right half of the matrix contains the Spearman correlation coeffients, while the italicised fields represent the corresponding p-values for each correlation score. Green highlighted fields indicate significant results at *p <* 0.1, *p <* 0.05, *p <* 0.01 with a darker green indicating a more significant result.

## Notes

### Competing Interest Statement

The authors have declared no competing interest.

