## Supplementary Material for "Incorporating uncertainty within dynamic interoceptive learning"

#### 1 BREATHING LEARNING TASK INSTRUCTIONS

BREATHING LEARNING TASK

***INSTRUCTIONS:***

In this task we will measure how you learn about your breathing. We will run a very short practice session first, and then we will run a longer version of the task (~30 min).

To run this task, we will ask you to breathe through a facemask while sitting at the computer. Most of the time it will be very easy to breathe just as you normally would.

During each trial, we will present one of two pictures on the screen. One of these symbols will mean that there is an 80% chance that it will become difficult to breathe for a short period (1-2 breaths), whilst the other symbol will mean that there will only be a 20% chance that it will become difficult to breathe. For each trial we will ask you to predict whether you think it will become difficult to breathe or not, according to the symbol. Following your prediction, a circle will appear on the screen, which is when this change in breathing either will or will not occur. Any resistance to breathing will only happen while you are inhaling – there will be no resistance as you breathe out – and breathing resistances will never be applied when there is no circle on the screen. After each breathing stimulus we will ask you to rate how difficult your breathing was. **NOTE:** The meaning of the symbols may **SWAP** at times during the experiment, but they are always paired together.

You will receive the key instructions again on the screen before you begin. Please feel free to ask any questions you may have now, or at any time during the practice session.

Please remember that you are able to stop at any time should you not wish to continue.

Thank you very much for participating!

**Figure S1.** Instructions for the Breathing Learning Task (BLT) provided to participants.

#### 2 PARAMETER RECOVERY ADDITIONAL RESULTS

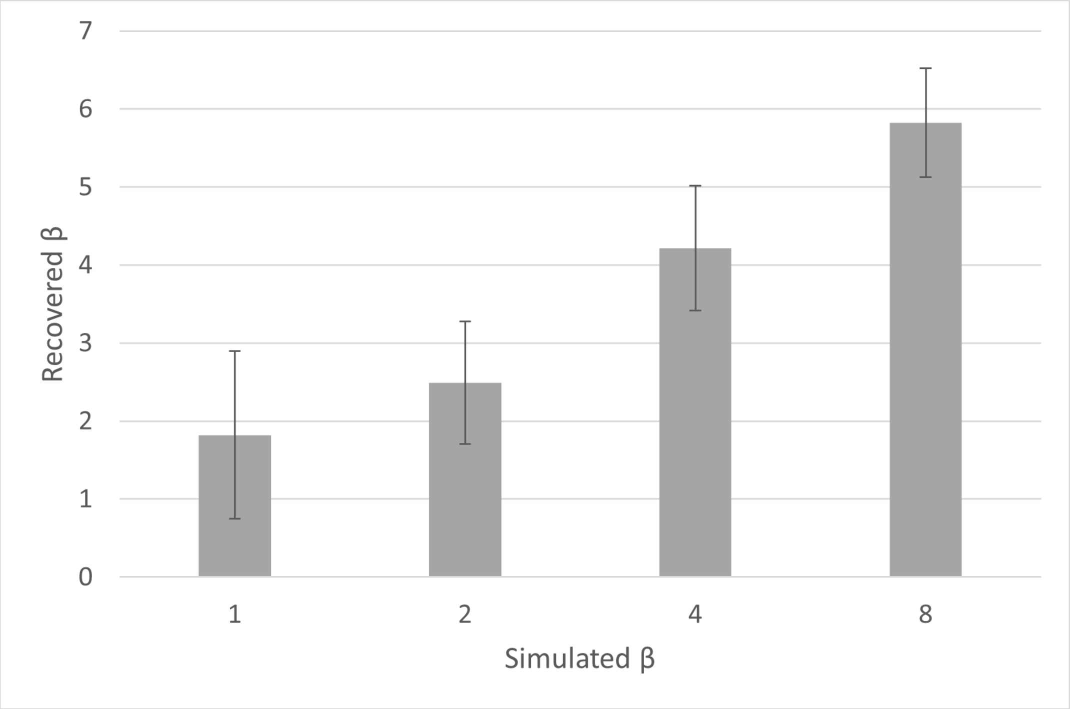


**Figure S2.** Parameter recovery for $\beta$ for the binary model, at four different simulated values of $\beta$. Error bars show the standard deviation of recovered values.

As seen in Figure S2, for simulations where $\beta = 1$, recovered $\beta$ values had a mean of 1.82 (SD = 1.08). At $\beta = 2$, recovered $\beta$ values had a mean of 2.49 (SD = 0.79), while at $\beta = 4$, recovered $\beta$ values had a mean of 4.22 (SD = 0.80). Finally, at $\beta = 8$, recovered $\beta$ values had a mean of 5.82 (SD = 0.70).

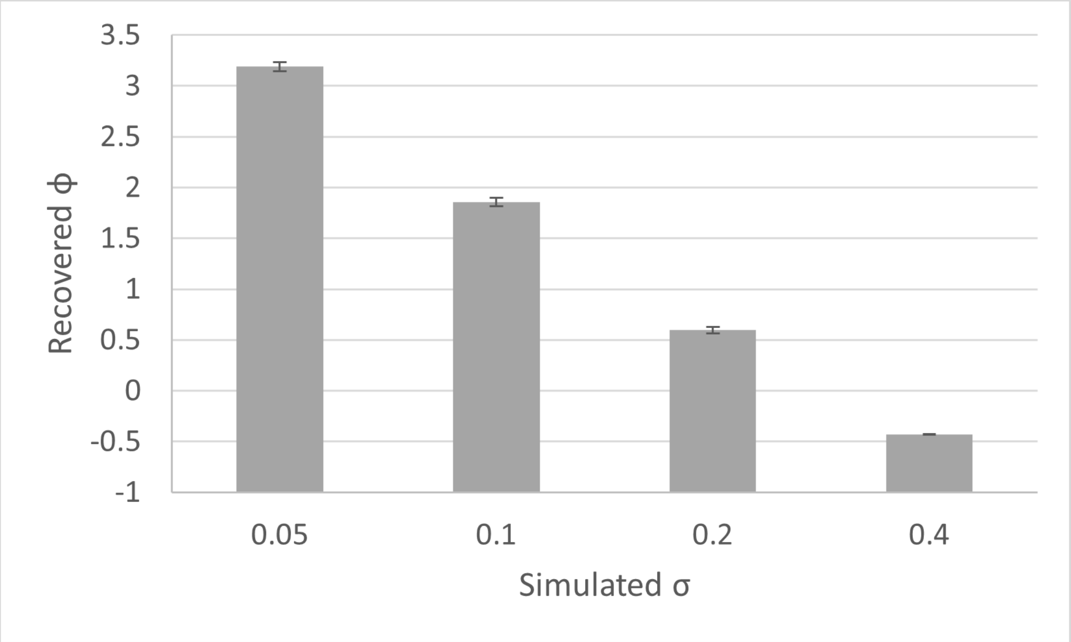


**Figure S3.** Relationship between $\sigma$ and $\phi$ from continuous model simulations, at four different simulated values of $\sigma$. Error bars show the standard deviation of recovered $\phi$ values.

As seen in Figure S3, $\sigma$ and $\phi$ values had an inverse relationship, with simulations at $\sigma = 0.05$ producing a mean $\phi$ of 3.19 (SD = 0.05), simulations at $\sigma = 0.1$ producing a mean $\phi$ of 1.86 (SD = 0.04), simulations at $\sigma = 0.2$ producing a mean $\phi$ of 0.60 (SD = 0.03), and simulations at $\sigma = 0.4$ producing a mean $\phi$ of -0.43 (SD = 0.005).

### 3 INDIVIDUAL FITS

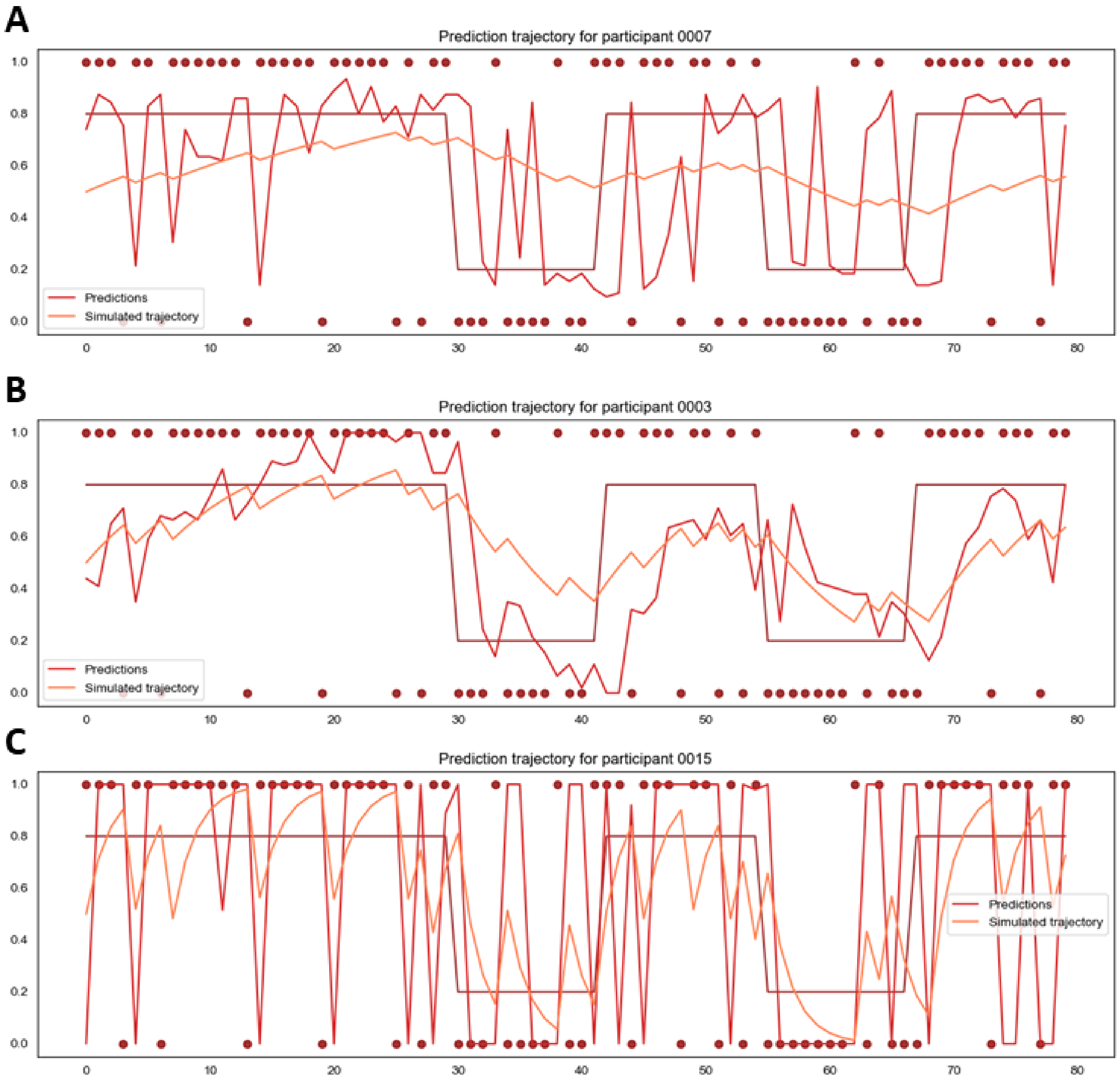


**Figure S4.** Individual fitted model trajectories (orange) compared to participant responses (bright red), with trial outcomes shown as dark red circles. **(A)** shows a participant with a low learning rate, **(B)** shows a participant with a moderate learning rate, and **C** shows a participant with a high learning rate as fitted by the continuous model.

### 4 DUAL LEARNING RATE MODEL RESULTS

#### 4.1 DLRM binary parameter recovery

Parameter recovery was successful at $\beta = 1$ ($r(\alpha_p) = 0.60$, $r(\alpha_n) = 0.62$), $\beta = 2$ ($r(\alpha_p) = 0.84$, $r(\alpha_n) = 0.85$), $\beta = 4$ ($r(\alpha_p) = 0.94$, $r(\alpha_n) = 0.94$), and $\beta = 8$ ($r(\alpha_p) = 0.95$, $r(\alpha_n) = 0.96$) for the binary model. Figure S5 shows a graphical representation of the dual learning rate binary model results, while Figure S6 shows a representation of the dual learning rate continuous model results.

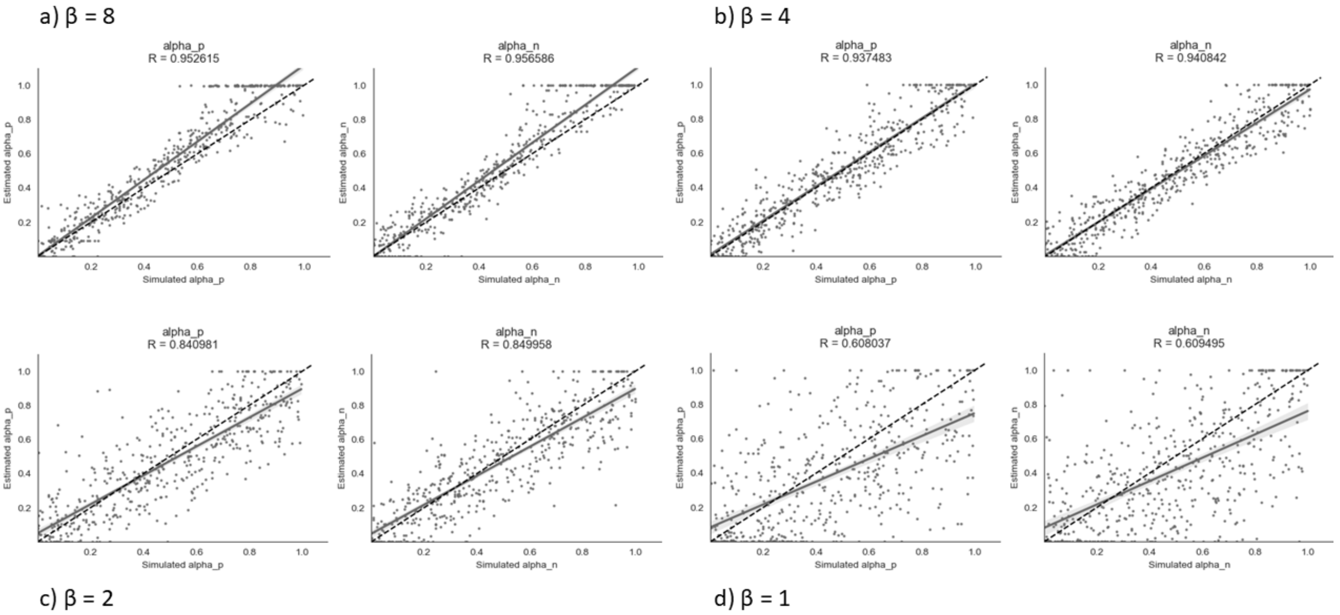


**Figure S5.** Parameter recovery of $\alpha_p$ and $\alpha_n$ for the dual learning rate binary model at different values of $\beta$.

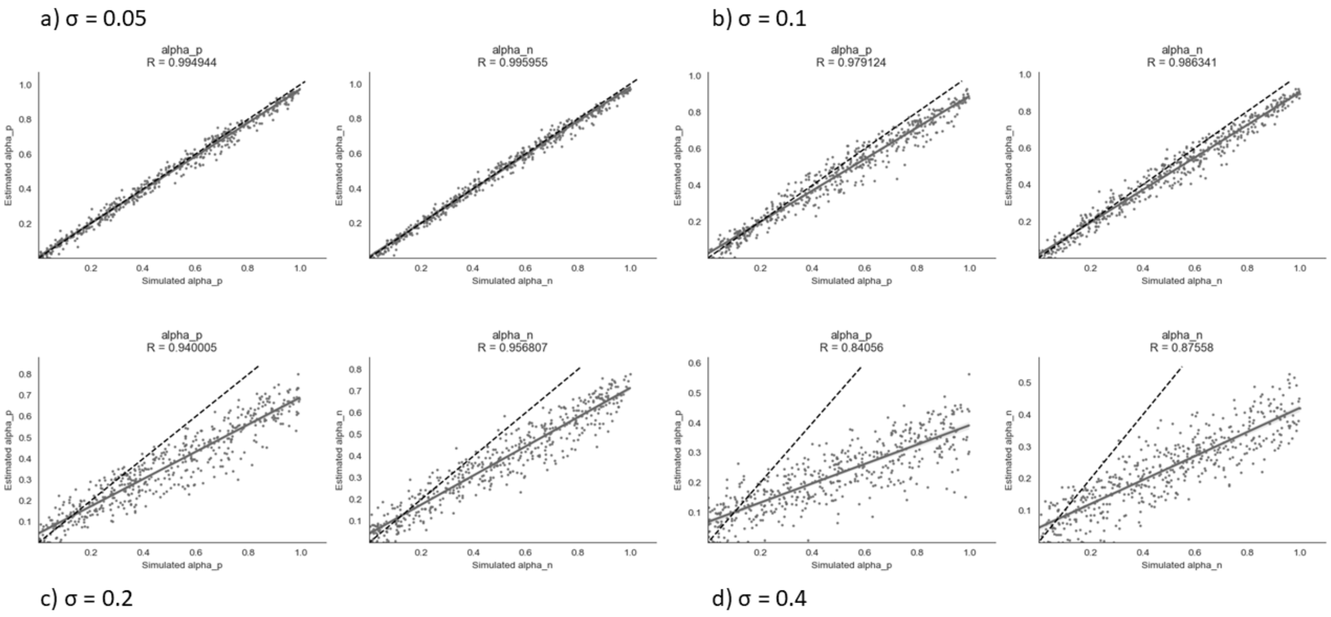


**Figure S6.** Parameter recovery of $\alpha_p$ and $\alpha_n$ for the dual learning rate continuous model at different values of $\sigma$.

#### 4.2 DLRM prediction trajectories

Figure S7 shows the prediction trajectories for the binary and continuous versions of the dual learning rate model.

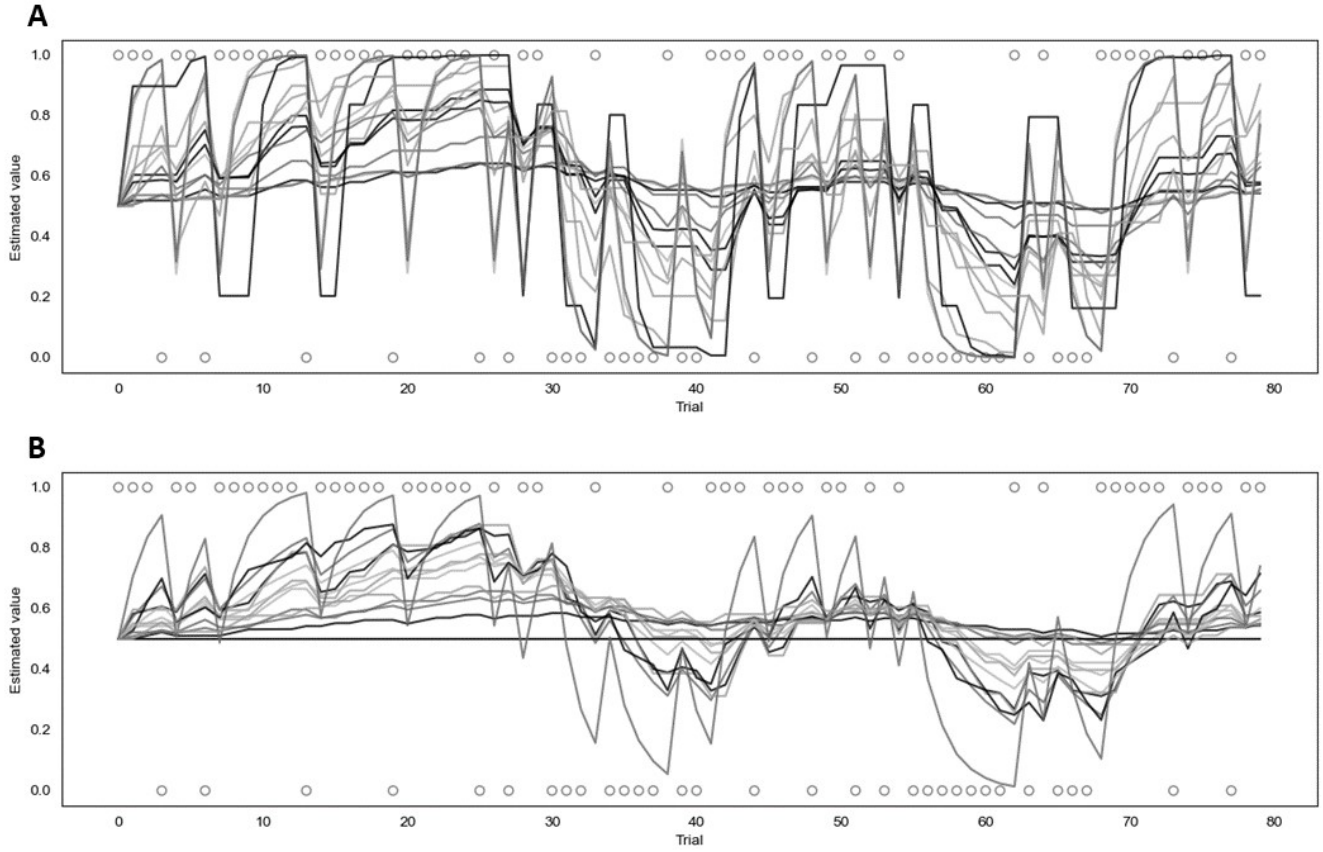


**Figure S7.** Fitted prediction trajectories for each participant for the dual learning rate model. **(A)** represents results from the binary version, **(B)** represents results from the continuous version.
